# BRD4 Deficiency Selectively Affects a Unique Developmental Subpopulation in Thymocytes

**DOI:** 10.1101/245662

**Authors:** Anne Gegonne, Qing-Rong Chen, Anup Dey, Ruth Etzensperger, Xuguang Tai, Alfred Singer, Daoud Meerzaman, Keiko Ozato, Dinah S. Singer

## Abstract

The bromodomain protein BRD4 is a driver in both inflammatory diseases and cancers. It has multiple functions, contributing to chromatin structure and transcription through its intrinsic HAT and kinase activities. Despite the wide-ranging characterization of BRD4, little is known about its in vivo function. In the present study, we have examined the role of BRD4 in T cell development by conditional deletion at various stages of thymocyte differentiation. We found that BRD4 is critical for normal T cell development. Surprisingly, BRD4 selectively regulates the progression of immature CD8 single positive (ISP) thymocytes into quiescent DP thymocytes. In striking contrast, BRD4 deletion does not affect the extensive proliferation associated with the differentiation of double negative (DN) into ISP cells. Nor does it affect the maturation of double positive (DP) into conventional CD4+ and CD8+ thymocytes. These studies lead to the unexpected conclusion that BRD4 selectively regulates preselection ISP thymocytes.

**On-line Summary:** Immature CD8 single-positive (ISP) thymocytes are identified as a molecularly-distinct thymocyte subpopulation, dependent on BRD4 for progression to the DP stage. DN and DP are BRD4-independent. These findings provide new insights into BRD4, a therapeutic target in inflammation and cancer.

## INTRODUCTION

Bromodomain protein 4 (BRD4) is a transcriptional and epigenetic regulator that plays a pivotal role in cancer and inflammatory diseases. First identified as a chromatin-binding factor that functions as a mitotic bookmark, it was subsequently shown to function as a scaffold that recruits the transcription elongation factor, P-TEFb, to sites of transcription and anchors super-enhancers (Dey et al., 2009; Jang et al., 2005; Loven et al., 2013; Yang et al., 2005). More recently, we have shown that BRD4 has both intrinsic histone acetyl transferase (HAT) and kinase activities (Devaiah et al., 2016a; Devaiah et al., 2012). BRD4 acetylation of nucleosomal histones – H3K122 in particular – at gene promoters it targets leads to octamer eviction and chromatin decompaction. This decompaction, in turn, allows the recruitment of the transcription preinitiation complex (PIC) and gene activation. BRD4 kinase activity phosphorylates the carboxy terminal domain (CTD) of RNA polymerase II (Pol II) at Ser2, which is necessary for pause release and productive transcription elongation. These activities thereby identify BRD4 as an active regulator that links chromatin structure and transcription.

BRD4 has been implicated in a broad spectrum of biological activities in a variety of normal and tumor cell types. Paradoxically, BRD4 has been shown to be necessary both to maintain stem cellness and for lineage-specific gene expression (Brown et al., 2014; Di Micco et al., 2014; Loven et al., 2013; Whyte et al., 2013; Wu et al., 2015). It is required for the in vitro differentiation of Th17 T cells, macrophage secretion of inflammatory cytokines and myogenic differentiation (Cheung et al., 2017; Roberts et al., 2017). In contrast, loss of BRD4 inhibits AML cell proliferation, driving their differentiation (Zuber et al., 2011). In many cell types, proliferation depends on BRD4 which functions throughout cell cycle: as a mitotic bookmark, at S (Dey et al., 2000; Dey et al., 2009; Mochizuki et al., 2008) and the G2/M transition through its interactions with various cell cycle factors (Farina et al., 2004). Deletion of BRD4 arrests cells at G1-S transition and is growth inhibitory to NIH3T3 and MEFs (Dey et al., 2009; Maruyama et al., 2002; Mochizuki et al., 2008)(unpublished data). In addition to regulating gene expression of both innate and adaptive immune cells (Bolden et al., 2014; Cheung et al., 2017; Dey et al., 2000; Mele et al., 2013; Schmidt et al., 2015), it regulates transcriptional events associated with mitosis, cell proliferation, viral latency and metabolism (Dey et al., 2009; Sakamaki et al., 2017; Tasdemir et al., 2016; You et al., 2004; You et al., 2009; You et al., 2005). Interestingly, although BRD4 is generally considered a transcriptional activator, it is a repressor of insulin secretion and metabolism (Barrow et al., 2016; Deeney et al., 2016). Alternative splicing of BRD4 gives rise to functionally distinct forms: a full length long form associated with tumor progression or a short form associated with metastasis (Alsarraj et al., 2013).

Despite the extensive characterization of BRD4 function in cell lines, little is known about its role in normal development beyond the fact that deletion is embryonic lethal (Houzelstein et al., 2002). The developmental program of T cells in the thymus is among the best characterized and provides an ideal in vivo system in which to assess the developmental role of BRD4 (Mingueneau et al., 2013; Rothenberg et al., 2016; Vacchio et al., 2016). The generation of T cells in the thymus results from the sequential differentiation of a series of thymocyte precursors. The earliest thymic immigrants from the bone marrow do not express any of the markers associated with mature T cells, namely the T cell receptor (TCR) and CD4/CD8 coreceptor molecules, and are called Double Negatives (DN). Within the thymus, the DN cells undergo a series of maturation steps punctuated by cycles of proliferation, rearrangement of the TCRβ gene and intracellular expression of the TCRβ protein. Further differentiation to the immature single positive (ISP) stage is accompanied by cell surface expression of CD8. The transition of the ISP to the double positive (DP), CD4+CD8+ thymocytes, requires a single round of cell cycle and is regulated by the transcription factors TCF-1, LEF-1 and RORγt, leading to surface expression of TCRαβ (Yu et al., 2004). DP thymocytes differentiate into either mature CD4+ or CD8+ single positive thymocytes that emigrate from the thymus to seed peripheral organs. DP thymocytes also give rise to iNKT cells; CD4+ SP thymocytes generate Foxp3+ Tregs.

The dependence on both robust proliferation and expression of lineage specific genes during thymic differentiation suggests that BRD4 may play a critical role during this process. The present study was undertaken to determine how BRD4 affects patterns of gene expression, proliferation and differentiation of thymocytes. By conditionally deleting BRD4 at various stages of thymic differentiation, we have established that BRD4 is not necessary either for proliferation at the DN stage or for the subsequent maturation of conventional CD4 and CD8 single positive thymocytes from the DP stage, although it is required for the generation of iNKT and Treg cells. Surprisingly, BRD4 selectively targets gene expression in the ISP cells: deletion of BRD4 in ISP down-regulates cell cycle and metabolic pathways, leading to a block in the transition to the DP stage. Furthermore, we unexpectedly identify the ISP as a cell type that is molecularly distinct from either the DN or DP subpopulations.

## RESULTS

### BRD4 Expression is Required Early in Thymocyte Development

BRD4 protein is expressed in the thymus and at similar levels across the different stages of thymocyte development. The levels of BRD4 in the thymus are comparable to those in peripheral CD4 and CD8 T cells and in B cells (Figure 1A, Figure S1A). To assess its role in thymocyte development, BRD4 was conditionally deleted at various stages of thymocyte development (Figures S2A and S2B, (Lee et al., 2001)). In thymi deleted of BRD4 by CD4-Cre, BRD4 protein levels were already reduced by 90% at the DP stage, allowing us to determine whether BRD4 affects the development of DP or CD4 and CD8 single positive thymocytes (Figure 1B). Despite the depletion of BRD4 protein, the total numbers and distribution of DN, DP, CD4+ and CD8+ thymocytes, as well as their surface expression of TCRβ and co-receptor molecules (Figure 1B), were comparable to those of the WT (Figure 1A). Thus, deletion of BRD4 by CD4-Cre did not significantly affect the differentiation of DP or their progression to single positive thymocytes. This leads to the conclusion that BRD4 is not essential to their development, despite the fact that it is expressed in those populations.

**Figure 1:**
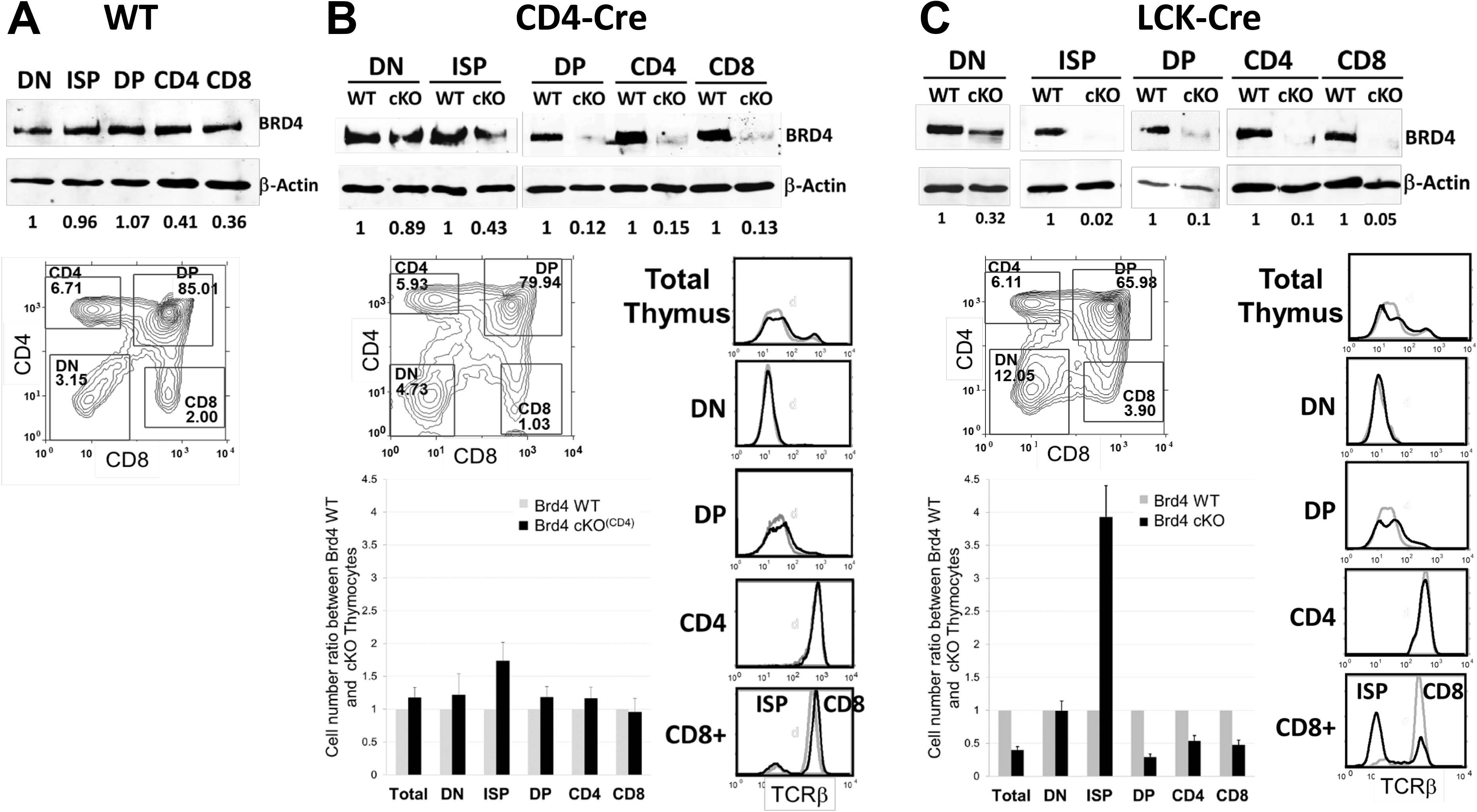
Brd4 deletion by LCK-Cre, but not CD4-Cre, affects thymocyte development. (A) Top: Immunoblot analysis of BRD4 and β-Actin in different populations of thymocyte. Level of BRD4 expression in thymocytes is normalized to β-actin and quantitated relative to DN. Bottom: Thymocytes from Brd4 wild type (WT) mice were analyzed by flow cytometry (FACS) based on their CD4 and CD8 cell surface expression to determine the distribution of DN, DP, and CD4 and CD8 SP cells. (B) Brd4 deletion by CD4-cre. Top: Immunoblot analysis of BRD4 and β-actin in different populations of WT or Brd4 F/- CD4-Cre+ (cKO) thymocytes. Level of BRD4 expression in thymocytes is normalized to β-Actin and quantitated relative to WT of each population. Bottom: Thymocytes from Brd4 F/- CD4-Cre mice (cKO^(CD4)^) were analyzed by FACS to determine the distribution of DN, DP, and CD4 and CD8 SP cells (left). Surface TCRβ for each cell population was determined in parallel (right). Cell number of each population was determined from the percentage and normalized relative to the WT cell numbers (Bar graph). The results represent mean+SEM of 5 independent analyses with 8 Brd4 F/- CD4-Cre+ mice, and 7 WT mice. (C) Brd4 deletion by LCK-cre.Top: Immunoblot analysis of BRD4 and β-actin in different populations of WT or Brd4 F/- LCK-Cre+ (cKO) thymocytes. Level of BRD4 expression in thymocytes is normalized to β-Actin and quantitated relative to WT of each population. Bottom: Thymocytes from Brd4 F/- LCK-Cre+ (cKO) mice were analyzed by FACS to determine the distribution of DN, DP, and CD4 and CD8 SP cells (left). Surface TCRβ for each cell population was determined in parallel (right). Cell number of each population was determined from the percentage and normalized relative to the WT cell numbers (Bar graph). The results represent mean+SEM of 13 independent analyses with 19 Brd4 F/- LCK-Cre+ mice, and 19 WT mice.

To further define the stage(s) of thymocyte development during which BRD4 expression is required, we deleted BRD4 with LCK-Cre (BRD4_cKO) which is expressed at an earlier stage than CD4-Cre (Figure S2B). Accordingly, BRD4 protein levels were markedly reduced in DN thymocytes in the BRD4_cKO thymus, relative to the wild type (WT) litter mates (Figure 1C) Deletion was complete by the immature CD8+ (ISP) stage; at subsequent stages of thymocyte development, BRD4 was virtually undetectable in the homozygous BRD4_cKO, relative to wild type (WT) (Figure 1C).

The early loss of BRD4 at the DN stage by LCK-Cre mediated deletion was accompanied by an aberrant distribution of thymocytes among the four subpopulations: whereas the percentage of DP was decreased, the percentages of DN and CD8^+^ thymocytes were markedly increased. (Figure 1C). The BRD4-deleted thymi were also significantly smaller than those of the wild type. A decrease of almost 65% in the DP population largely accounts for the overall reduction in thymus size (Figure 1C, bottom). The numbers of CD4+ and CD8+ thymocytes were also proportionally reduced (Figure 1C, bottom).

In contrast to the aberrant patterns of co-receptor expression, TCRβ expression in the LCK-cre BRD4-depleted thymus paralleled that of the wild type in all subpopulations, with one notable exception (Figure 1C). Within the CD8+ population, a large sub-population of the BRD4-deleted cells did not express surface TCRβ. The increase in the proportion of CD8+ thymocytes that lack surface TCRβ expression indicated an accumulation of immature single positive (ISP) thymocytes which are the developmental intermediate stage between DN and DP. They are characterized by the expression of CD8, but not TCR, on their cell surface (Takahama et al., 1992). Taken together, these findings identify a block in differentiation at the ISP stage in the absence of BRD4 which resulted in an increased proportion of ISPs and a failure of normal developmental processes.

Deletion of BRD4 by VAV-Cre, which is first expressed in hematopoietic stem cells (HSC) in the bone marrow (Figure S2B and (Georgiades et al., 2002)) was embryonic lethal, with only 2 live births of more than 160 total (Dey in preparation). The two pups that survived had few thymocytes, among multiple other defects in hematopoiesis (Dey, in preparation), consistent with the previous observation that adult bone marrow depletion of BRD4 by shRNA suppresses hematopoiesis (Bolden et al., 2014).

Taken together, these results lead to the surprising conclusion that BRD4 is required during the early stage(s) of thymic development, but is not required for the maturation of phenotypically normal thymocytes at the DP or conventional SP stages. Thus, BRD4 is critical during a narrow window of thymic differentiation prior to the DP stage and focused on the ISP stage.

### BRD4 deletion blocks thymocyte development at the ISP stage

The increased fraction of TCRβ-negative CD8+ thymocytes in the thymus deleted of BRD4 by LCK-cre (Figure 1C) indicated an accumulation of ISP. To directly examine the ISP populations in wild type and BRD4-deficient thymus, we determined the fraction of CD8+ ISP cells within the total surface TCRβ-negative thymocyte population (Figure 2A). The deletion of BRD4 (BRD4 F/- LCK-Cre+) was accompanied by a large increase in TCRβ-negative cells in the thymus, relative to the WT (Figure 2A, left). As shown in Figure 2A, among the total TCR β-negative cells in the thymus (gated to the left of the vertical line in Figure 2A, left), the percentage of CD8+ ISP cells increased nearly 20-fold to 4.8% in the BRD4-deleted thymus, compared with 0.26% in the WT (Fig. 2A, middle and right panels. Similarly, within the total thymocyte population, the WT consisted of 0.16 % ISP, whereas the ISP population in the BRD4-deleted thymus increased by 10-fold to 1.6 %, consistent with a block in differentiation (data not shown). This increased percentage of ISP cells led to a correspondingly higher absolute number of ISP in the BRD4-deficient thymus (Figure 2B).

**Figure 2:**
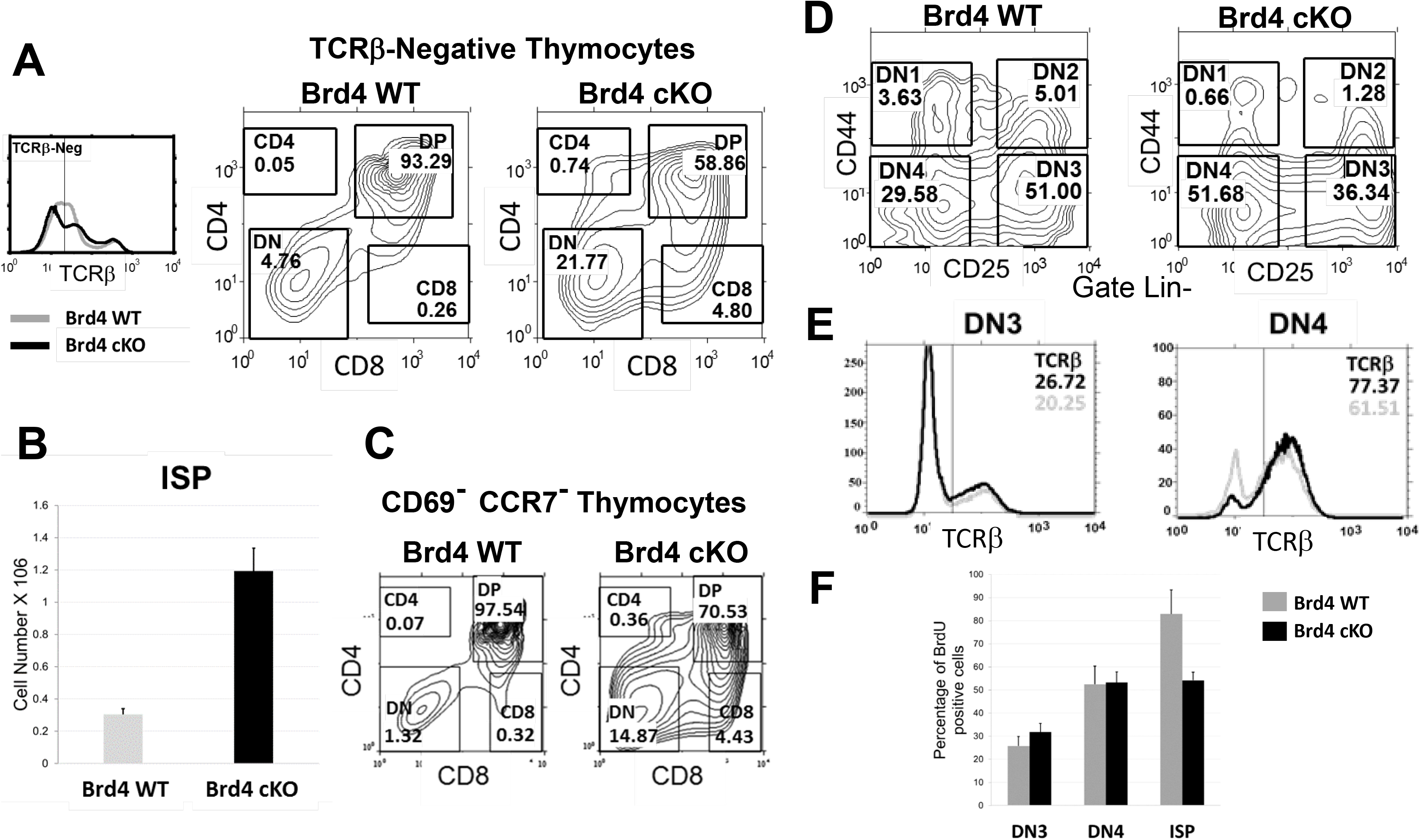
BRD4 regulates the transition from ISP to DP. (A) Thymocytes from Brd4 wild type (WT, gray line) and Brd4 F/- LCK-Cre mice (cKO, black line) were analyzed by FACS for surface TCRβ expression (left panel). Cells with low surface TCRβ (left of vertical line) were analyzed by FACS for CD4 and CD8 cell surface expression. ISP were identified as CD8 positive cells with no TCRβ surface expression. BRD4 WT, middle panel; Brd4 F/- LCK-Cre+ (cKO), right panel. (B) Quantification of ISP in thymi from Brd4 wild type (WT) and Brd4 F/- LCK-Cre (cKO). The results represent mean+SEM of 13 independent analyses with 19 Brd4 cKO mice and 19 WT mice. (C) CD69^low^, CCR7^low^ thymocytes from Brd4 wild type (WT) and Brd4 cKO mice were analyzed for the expression of CD4 and CD8. The data are representative of 3 independent experiments. (D) DN thymocytes WT or cKO, defined as negative for lineage commitment markers (Lin-), were categorized into the 4 stages of differentiation based on cell surface CD25 and CD44 markers as described in Methods. The data are representative of 4 independent experiments. (E) Intracellular TCRβ expression in DN3 and DN4 thymocytes from WT (gray line) and Brd4 cKO (black line) mice. DN3 (left panel) and DN4 (right panel) thymocytes were stained as described in Methods, fixed and cytoplasmic TCRβ expression analyzed by FACS. The data are representative of 2 independent experiments. (F) BrdU incorporation by DN3, DN4 and ISP thymocytes from WT and Brd4 cKO. The 3 cell types were analyzed for BrdU incorporation 4 hours after injection. Data shown summarize three independent experiments for DN and 2 for ISP and represent mean+SEM.

The block in the transition from ISP to DP in the BRD4-deficient thymus was further documented by analyzing the distribution of the immature pre-selection thymocyte subsets. Pre-selection thymocytes are characterized by their CD69^low^ and CCR7^low^ phenotype. In WT mice, these pre-selection CD69^low^CCR7^low^ thymocytes consisted primarily of DP thymocytes (97.5%), with a small fraction of DN (1.3%) and ISP (0.32%) cells (Figure 2C). In striking contrast, in the BRD4-deficient thymus, the percentage of CD8+ (ISP) cells increased sharply, representing 4.4% of the CD69^low^CCR7^low^ population, about a 14-fold increase relative to the wild type (Figure 2C). The identity of the CD8+ cells as ISP was confirmed by the demonstration that they express CD8αβ, CD24 and no surface TCRγδ (Figure S3, data not shown). Thus, loss of BRD4 leads to an accumulation of ISP cells, reflecting a block in the transition from ISP to DP thymocytes.

We next examined the possibility that BRD4 also regulates DN maturation. The four DN subpopulations, DN1–4, are distinguished by their expression of the cell surface markers CD25 and CD44. Deletion of BRD4 by LCK-Cre resulted in a shift in the distribution of DN cells, with an increase in the percentage of the more differentiated DN3 and DN4 (Figure 2D). A decrease in the size of DN4 thymocytes also accompanied the loss of BRD4 (Figure S4). Although modest, these changes in DN cells in the absence of BRD4 led us to examine whether their maturation was defective.

Maturation from DN cells to ISP cells requires both the rearrangement of the TCRβ genes and extensive proliferation (Rothenberg et al., 2016; Takahama et al., 1992). A defect in either TCRβ rearrangement or in proliferation might result in a failure of BRD4 deleted DN cells to differentiate, leading to their accumulation. We first examined the extent of TCRβ rearrangement, as assessed by internal staining of DN3 and DN4 cells. No differences were detected in the level of intracellular TCRβ between the BRD4-deficient and wild type thymocytes, indicating that BRD4 is not required for the rearrangement or expression of TCRβ (Figure 2E). Therefore, the block in differentiation resulting from BRD4 deficiency occurs subsequent to, and is independent of, TCRβ rearrangement.

To determine whether BRD4 deletion affects the proliferation of DN thymocytes, we examined their ability to incorporate BrdU. Mice were injected peritoneally with BrdU and their DN thymocytes analyzed by flow cytometry for BrdU incorporation 4 hours post injection (Figure 2F). Surprisingly, the extent of BrdU incorporation was indistinguishable between the BRD4-deleted and WT thymocytes at either DN3 or DN4 stages, despite the fact that BRD4 levels were already reduced by two-thirds in total BRD4 deleted (BRD4F/-LCK-Cre+) DN thymocytes (Figure 1C). Thus, BRD4 does not appear to be required for proliferation of DN thymocytes (Figure 2F). In contrast, the ISP cells incorporated significantly less BrdU, suggesting a defect in proliferation at the ISP stage (Figure 2F).

The failure to detect defects in either TCRβ rearrangement or proliferation in BRD4-deficient DN3 or DN4 cells indicates that deletion of BRD4 at the DN2/3 stage does not impair maturation of DN thymocytes through to the DN4 stage. Thus, BRD4 is not required for the differentiation of DN thymocytes. Rather, a major role of BRD4 in thymocyte development is in regulating the transition from ISP to DP.

### ISP Thymocytes are a Distinct Thymocyte Subpopulation

To begin to understand the molecular basis of the block in differentiation in thymocytes deleted of BRD4 by LCK-Cre, we first examined the gene expression profiles of DN, ISP, and DP thymocytes from wild type thymus by RNA-seq to determine whether the ISP are simply a transitional cell or a functionally distinct subpopulation. As expected, each of the transitions from one developmental stage to the next was accompanied by large changes in gene expression (Figure 3A). Surprisingly, the greatest differences in gene expression in the WT were associated with the ISP subpopulation in its transition to the DP stage (Table 1; Figure 3A). Furthermore, 744 genes (7.5% of the total) were differentially expressed in ISPs relative to both the DN and the DP subpopulations (Figures 3B and 3C).

**Figure 3:**
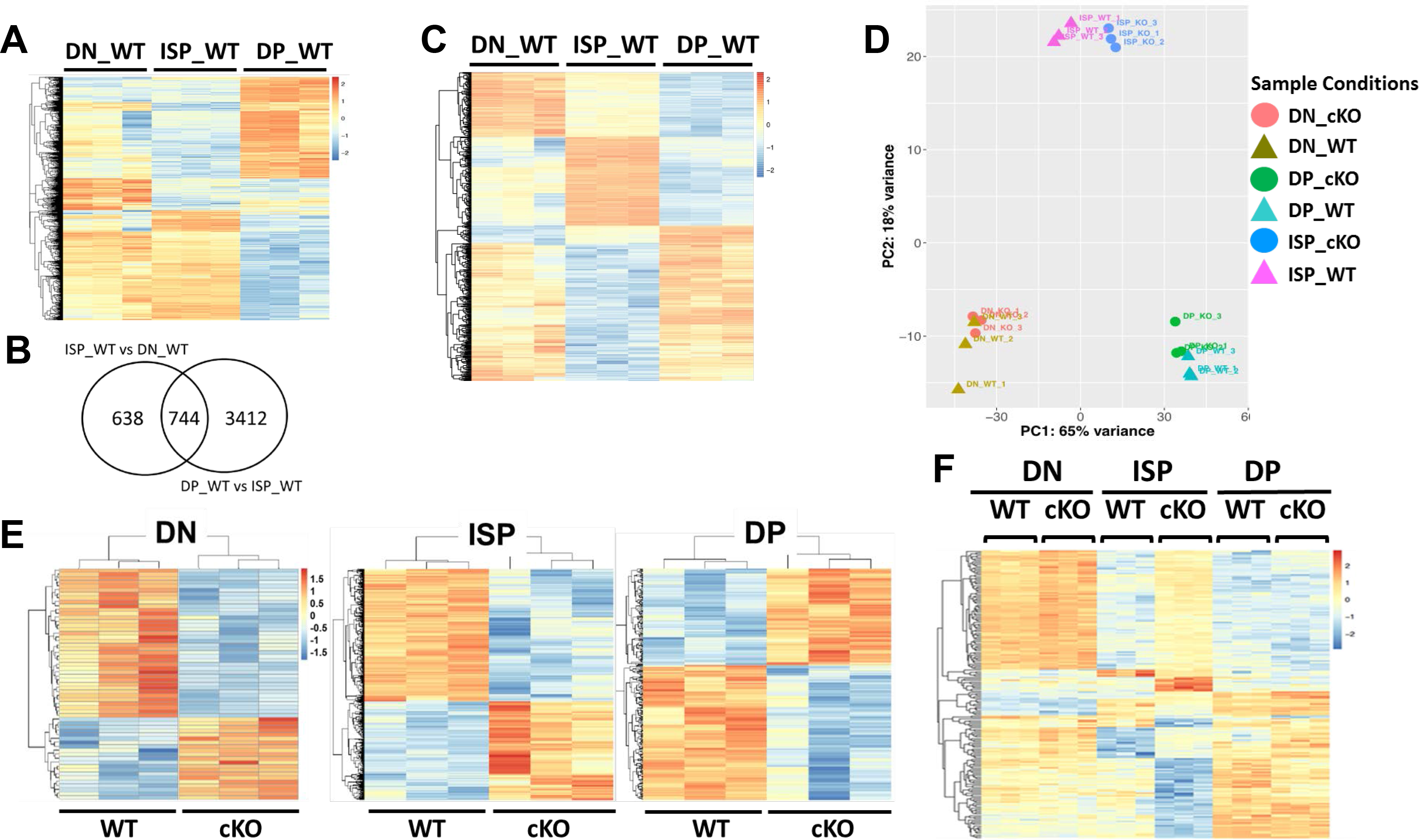
Brd4 deletion preferentially targets ISP thymocytes. (A) Heat map comparing RNA expression profiles of DN, ISP and DP thymocytes from WT thymus. (B) Venn diagram summarizing the differences in gene expression between WT ISP thymocytes and WT DN and DP, respectively. (C) Heatmap of the 744 genes differentially expressed in WT ISP relative to DN and DP. (D) Principal components analysis (PCA) of gene expression profiles of DN, ISP and DP thymocytes from WT and BRD4 conditionally deleted thymocytes (cKO). (E) Heat maps illustrating the effects of BRD4 deletion on gene expression in DN, ISP and DP thymocytes. The heatmaps represent genes with a signal intensity fold change >2 and FDR <0.01 in BRD4 thymocytes conditionally deleted by LCK-Cre (cKO) relative to WT for each cell type as indicated. (F) Heat map comparing the expression of the 194 BRD4-dependent genes of the 744 that are differentially expressed genes in WT ISP relative to DN and DP.

**Table 1.** Differential Gene Expression in Thymocyte Subpopulations and in the Absence of BRD4.

| **Comparison** | **Genes Differentially Expressed (#)** | **Up-Regulated (#)** | **Down-Regulated in (#)** |
| --- | --- | --- | --- |
| Wild Type |  |  |  |
| ISP_WT vs DN_WT | 1382 | 350 | 1032 |
| ISP_WT vs DP_WT | 4156 | 2143 | 2013 |
| CD4_WT vs DP_WT | 3127 | 1778 | 1349 |
| CD8_WT vs DP_WT | 3223 | 1811 | 1412 |
| CD4_WT vs CD8_WT | 842 | 462 | 380 |
| **Differential Expression in vivo** |  | **Up-Regulated in BRD4_cKO(#)** | **Down-Regulated in BRD4_cKO (#)** |
| DN_cKO vs DN_WT | 107 | 35 | 72 |
| ISP_cKO vs ISP_WT | 1168 | 517 | 651 |
| DP_cKO vs DP_WT | 349 | 166 | 183 |
| CD4_cKO vs CD4_WT | 388 | 53 | 335 |
| CD8_cKO vs CD8_WT | 308 | 64 | 244 |
| **Differential Expression in Culture** |  |  |  |
| cult.DP_WT vs Input_WT | 3194 |  |  |
| cult.ISP_cKO vs Input_cKO | 1902 |  |  |
| cult.DP_cKO vs cult.ISP_cKO | 201 |  |  |
| Cult.DP_cKO vs cult.DP_WT | 299 |  |  |

Legend for Table 1: Differentially expressed genes were defined as >2-fold difference in signal intensity, with FDR of <0.01.

Pathway analyses revealed that the gene sets that are differentially expressed in the wild type ISP population compared to both WT DN and DP are primarily associated with cell cycle and metabolic pathways (Figure S5A). In contrast, most of the changes in gene expression between DN and ISP were clustered in immune and T cell development pathways, although they also included cell cycle pathways (Figure S5B). Consistent with the known quiescence of DP cells, the transition from ISP to DP cells was accompanied by a down-regulation of cell cycle pathways, as well as an up-regulation of cell death and T cell development pathways (Figure S5C).

Principal components (PCA) analysis highlighted the fact that the ISP population is distinct and no more related to either DN or DP than they are to each other (Figure 3D). These findings document the ISP as a distinct cell type whose gene expression profile is not simply a hybrid intermediate state between DN and DP. Rather, the pattern of gene expression in ISPs is unique and not found in either DN or DP and may be required to support the transition to DP.

### BRD4 Targets a Set of ISP-Specific Transcripts

We next examined by RNA-seq the effect on gene expression of BRD4 deletion in DN, ISP and DP thymocyte subpopulations (Figure 3E). Deletion of BRD4 did not reduce gene expression globally: All of the subpopulations, in both WT and BRD4 cKO expressed approximately the same total number of genes. However, in allof the subpopulations, BRD4 deletion selectively affected levels of gene expression (Table 1). Pathway analysis revealed that many genes associated with immune cell function are differentially expressed in all thymocyte subpopulations as a result of BRD4 deletion (Table 2).

**Table 2.**
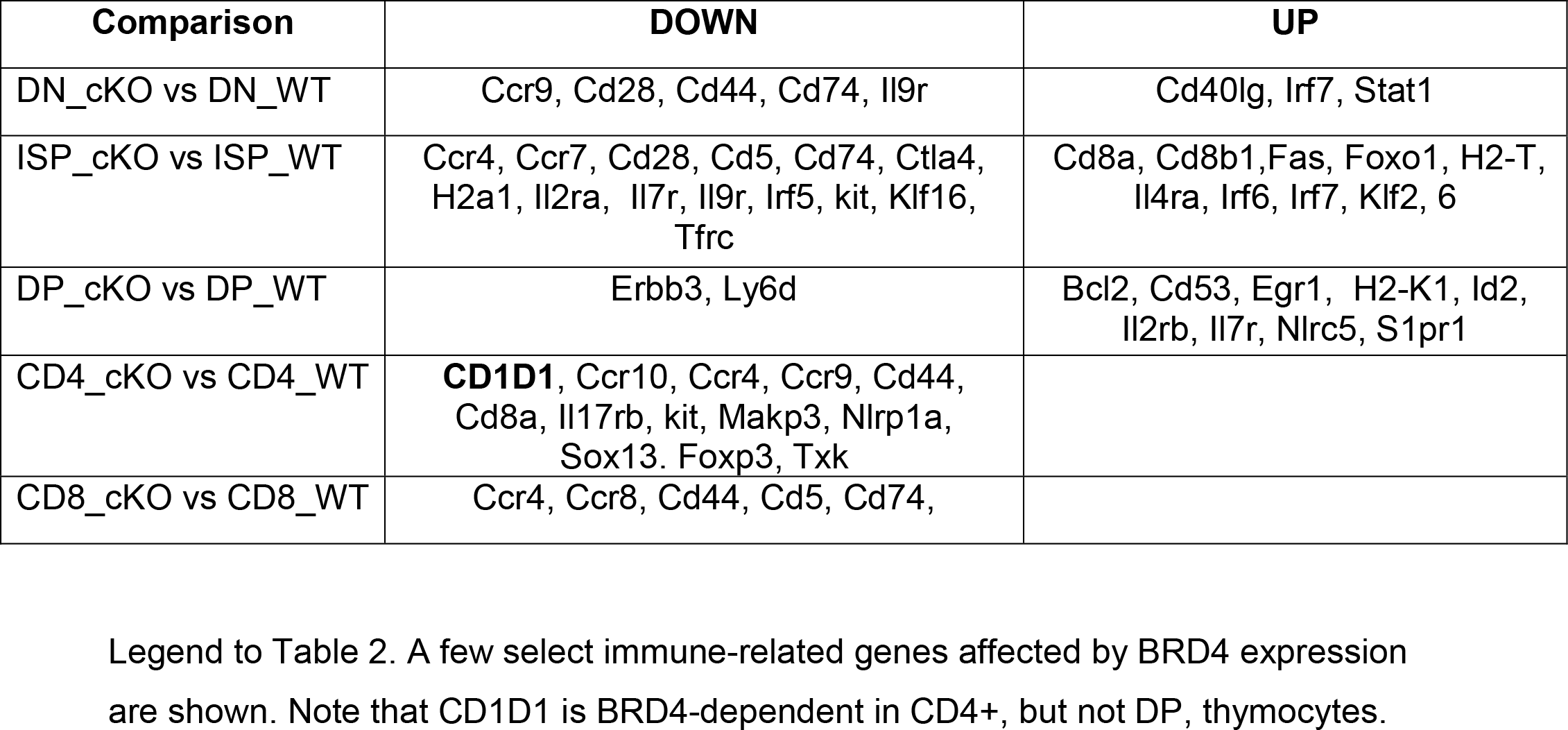
Examples of immune-related genes regulated by BRD4.

The largest effect of BRD4 deletion on gene expression was observed in the ISP cells (Figure 3E, Table 1). Strikingly, 1168 genes were differentially expressed between the BRD4 deficient and WT ISP cells: 517 were increased and 651 decreased in expression. Of those, 194 were among those differentially expressed in ISP relative to both DN or DP (Figure 3F). Among the pathways most affected were those related to metabolism and cell growth, as well as immune signaling pathways (Figure S6A), which were all down-regulated in the absence of BRD4. Thus, BRD4 primarily functions as an activator of the major ISP pathways. Importantly, these results indicate that the ISPs are a distinct thymocyte subpopulation that is selectively regulated by BRD4.

Although we detected no significant phenotypic or developmental defects in either the DN or DP subpopulations in the absence of BRD4, we did observe modest differential gene expression in both subpopulations. Deletion of BRD4 resulted in the differential expression of only 107 genes in the DN population, despite the nearly two-thirds loss of BRD4. Expression was increased in 35 transcripts and decreased in 72 (Table 1). These changes may account for the small differences observed in the size of DN and the DN4/3ratio. Similarly, a relatively modest number of genes was differentially expressed in BRD4_cKO DP cells compared to WT (Table 1). Among the genes differentially expressed in the DP thymocytes, about half were up-regulated in the absence of BRD4, identifying BRD4 as a repressor of those genes. Indeed, the overall effect of BRD4 is as repressor of the major DN and DP immune and signaling pathways (Figure S6B).

Despite the alterations in gene expression resulting from BRD4 deletion, principal components and t-SNE analyses highlighted the fact that the differences among the DN, ISP and DP subpopulations are notably greater than the differences between the WT and BRD4_cKO within a subpopulation (Figure 3D and Figure 6E).

### BRD4 Regulates Glycolytic and Myc Pathways in ISP Thymocytes

RNA-seq analysis indicated that deletion of BRD4 affected not only immune pathways, but also profoundly affected glycolytic pathways in ISP thymocytes (Figure 4; Figure S6A). Among the genes differentially expressed in ISP following BRD4 depletion were glucose transporters, leading to the prediction that BRD4_cKO ISP cells should be defective in glycolysis. To test this prediction, we compared the ability of BRD4_cKO ISP thymocytes to take up the glucose analog, 2-NBDG (Figure 4A, top). As predicted, BRD4 cKO ISP thymocytes were impaired in glucose uptake, relative to the WT (Figure 4A, top). In contrast, BRD4 deletion did not affect glucose uptake in DP (Figure 4A, top). Although not predicted by the pathway analysis, glucose uptake in DN thymocytes was slightly reduced in the absence of BRD4. Mitochondrial pathways were not identified in the analyses; accordingly, mitochondrial mass was not detectably affected by the absence of BRD4 in any of the thymocyte subpopulations (Figure 4A, bottom).

**Figure 4:**
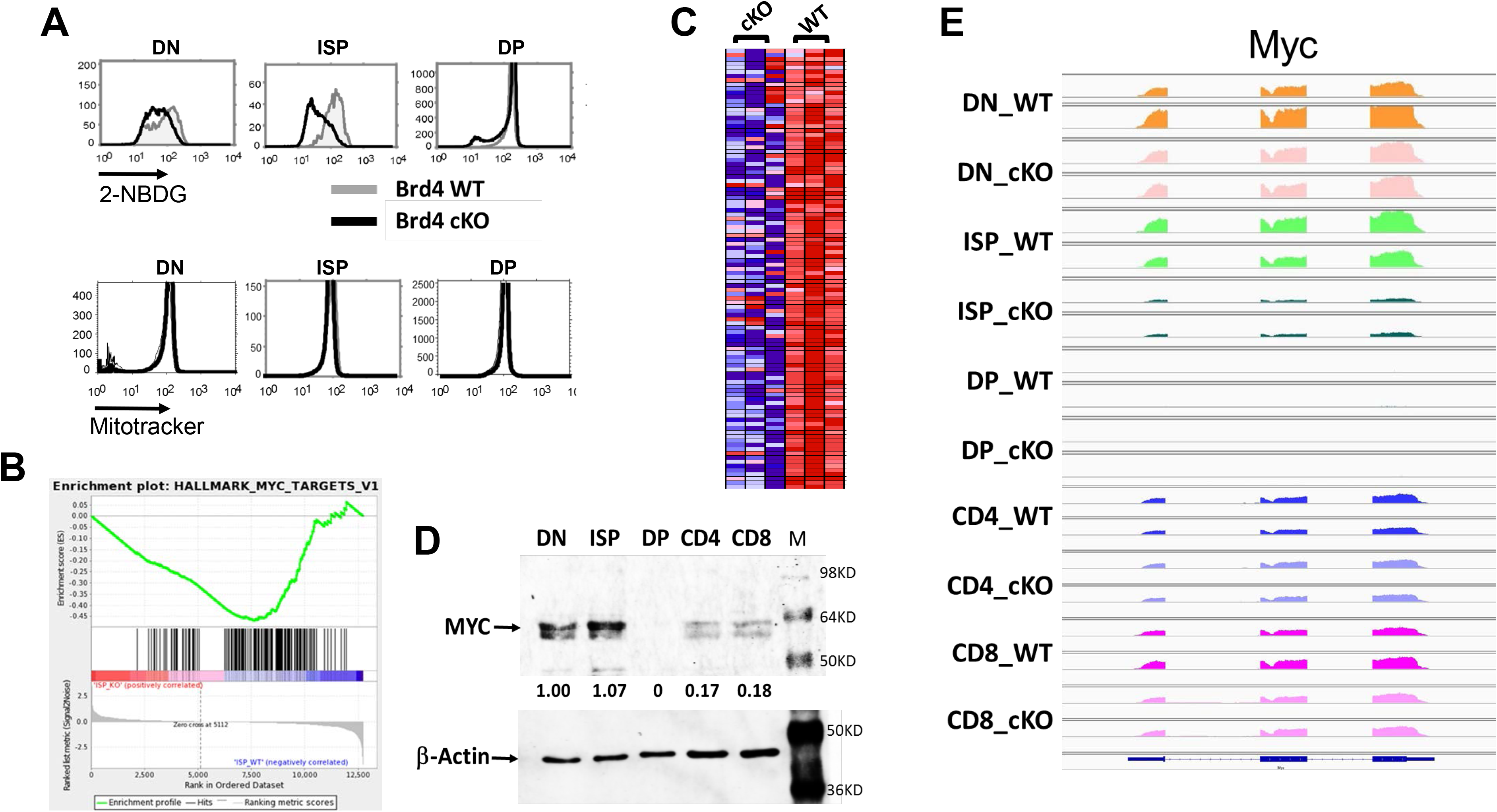
BRD4 regulates glycolytic and myc pathways in ISP thymocytes. (A) Top: Glucose uptake in WT and BRD4_cKO DN, ISP and DP thymocytes, as assessed by uptake of the Glucose analog 2_NBDG and assessed by FACS. WT, gray; BRD4_cKO, black. Bottom: Mitochondrial mass of Brd4 WT and cKO DN, ISP and DP thymocytes, as assessed by mitotracker uptake and FACS. (B) Enrichment plot of the c-Myc target gene set in ISPs. The c-Myc_targets_v1 gene set is significantly down-regulated in the Brd4_cKO (p<0.0001). The green curve shows the running sum of enrichment score (ES) for ranked genes. The hashmarks under the plot represent the genes that are the leading edge subset which accounts for the gene set’s enrichment signal. (C) Heat map showing the normalized expression of top down-regulated myc_target_v1 genes in BRD4_cKO ISP thymocytes. (D) Immunoblot analysis of MYC and β-actin in different populations of thymocyte. (E) Browser views of Myc gene expression in WT DN, ISP, DP, CD4 and CD8 thymocytes.

BRD4 deletion in ISP cells also significantly repressed many myc target pathways but increased expression of other genes, notably inhibitors of cell cycle progression such as Cdkn1b and 2d. (Figures 4B and 4C).To characterize the role of BRD4 in regulating Myc target pathways, we examined Myc RNA and proteins levels in the individual wild type thymocyte subpopulations (Figures 4D and 4E). Myc RNA was robustly expressed in all subpopulations except for the DPs which express no detectable Myc consistent with their quiescent phenotype and small size (Figure S4). Accordingly, MYC protein levels are highest in the ISPs and DNs, somewhat lower in CD4+ and CD8+ thymocytes and undetectable in DPs (Figure 4D).

Although myc was expressed in all thymocyte subpopulations except DP, BRD4 deletion only eliminated Myc RNA expression in the ISP thymocytes (Figure 4E). It did not affect Myc RNA levels in the DNs, consistent with our *in vivo* observations that deletion of BRD4 did not abrogate the proliferation of DN thymocytes (Figure 2F). Myc expression in CD4+ or CD8+ SPs was similarly unaffected (Figure 4E). Thus, surprisingly, myc expression is only BRD4-dependent at the ISP stage.

Taken together, these findings validate the predictions of the RNA-seq data, indicating that the RNA-seq data provide an accurate assessment of the molecular phenotypes of the thymocyte subpopulations. Importantly, the RNA-seq and functional data establish ISP thymocytes as a distinct subpopulation, not a hybrid transitional state, that is selectively regulated by BRD4.

### BRD4-Deficient ISP Fail to Mature to DP During In Vitro Culture

The RNA-seq data provided a molecular definition of each of the thymocyte subpopulations. However, they did not define the dynamic changes in gene expression that accompany the transition from ISP to DP. Maturation of ISP to DP requires a single round of cell cycle (Yu et al., 2004) and can be reproduced in an in vitro culture, allowing us to identify those genes dynamically associated with differentiation. Following an overnight culture, WT ISP doubled in number and all matured to DP thymocytes (Figure 5A, B). In contrast, BRD4-deficient ISP did not divide and were severely impaired in their ability to differentiate into DP cells. Less than half of the BRD4_cKO ISP expressed CD4 after the overnight culture and at levels lower than the WT CD4+ cells (Figures 5A and 5B). However, those BRD4-deficient ISP that expressed CD4+ following the *in vitro* culture expressed surface TCR at levels comparable to the WT, indicating that BRD4 is not necessary for surface TCR expression (Fig 5C, bottom). Further, our results lead to the surprising conclusion that although BRD4 is not required for DN proliferation, it is necessary for the single round of cell cycle that leads to the transition from ISP to DP. This raises the intriguing possibility that the requirements for cell division in these populations are distinct.

**Figure 5:**
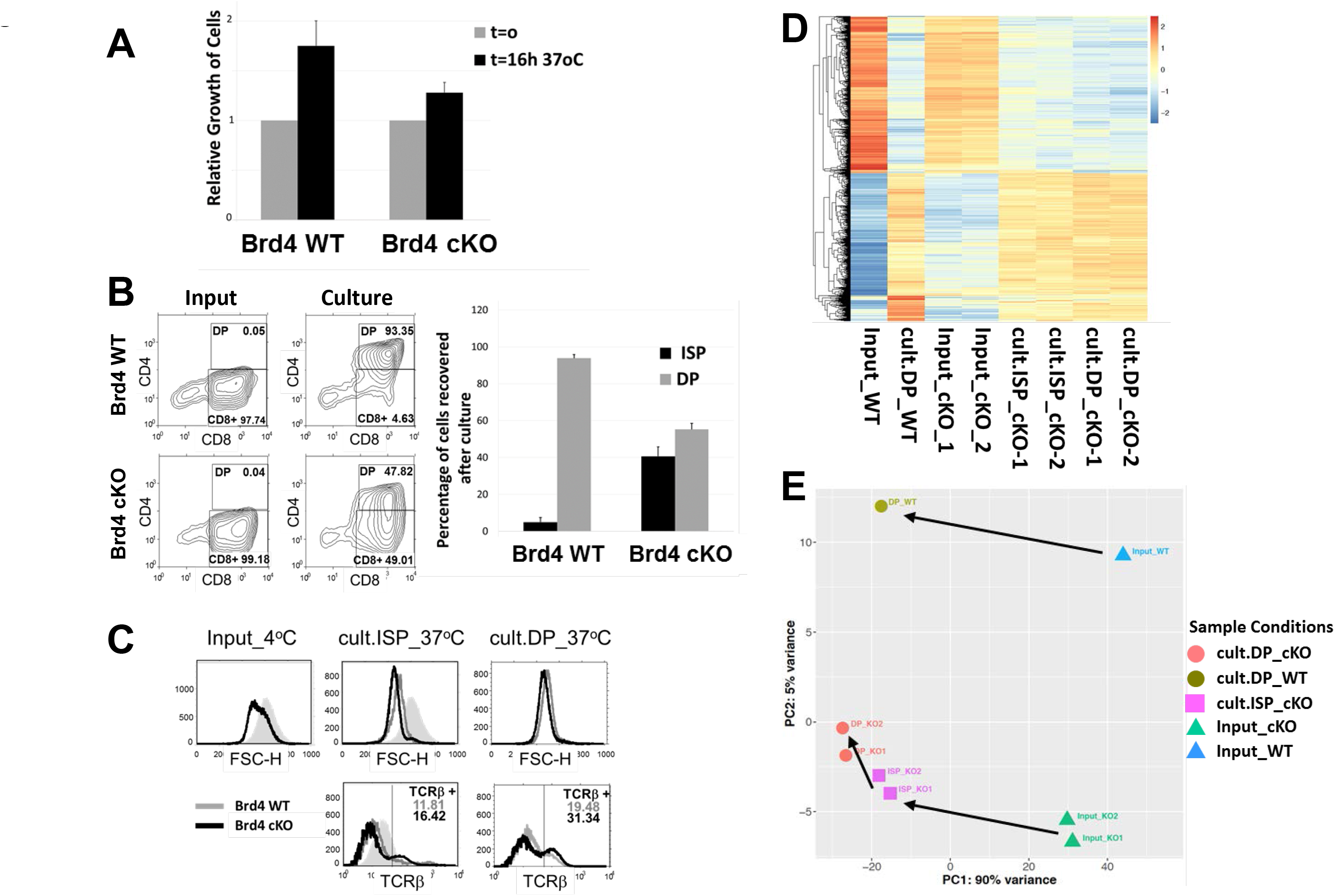
BRD4_cKO ISP are blocked in their maturation into double positive thymocytes in vitro. (A) ISP from BRD4 cKO and WT thymi were cultured overnight and the extent of their proliferation was quantified as cell recovery (percent of input) after the overnight culture; data are the mean+SEM of 4 independent experiments. (B) Extent of maturation into DP (culture panel), relative to the input (input panels), of ISP from WT (top) and cKO (bottom panel) was assessed by flow cytometric measurement of CD4 and CD8 surface marker expression. The histogram (right) represents the mean+SEM of 4 independent cultures of Brd4 cKO ISP and 3 WT ISP. (C) FACS of the input ISP remaining after culture (cult.ISP_37^o^C) and DP appearing after culture (cult.DP_37^o^C) (from B) assessing their size by forward scatter (top) and TCRβ surface expression (bottom). WT, gray; Brd4 cKO, black; shaded peak, WT ISP input. The data are representative of 4 independent experiments. (D) The heat map was plotted based on 3194 genes that are differentially expressed between DP_WT after overnight culture (cult.DP_ WT) and input ISP_WT (Input_ WT). Input ISP from Brd4 cKO mice (Input_cKO), remaining cKO ISP (cult.ISP_cKO) and cKO DP (cult.DP_cKO) after overnight culture. (E) PCA representation of the WT and cKO input ISP and thymocyte populations after overnight culture. 38

During normal development in the WT thymus, the transition to the DP stage is accompanied by a reduction in the size of the thymocytes. The BRD4-deficient ISP harvested from the thymus were smaller than WT ISP, presumably reflecting the loss of Myc (Figure 5C, top). Nevertheless, the ISP that progressed to the CD4+ stage in culture, as well as those that did not, underwent the further reduction in size associated with the normal transition to DP (Figure5C, top). From these data, we conclude that although BRD4-deficient ISP undergo some of the phenotypic changes associated with differentiation – indicative of signaling – they are severely restricted in their ability to mature to the DP stage, which is manifested by an impaired ability to undergo cell division and reflected in the accumulation of ISPs in the BRD4-deficient thymus.

To characterize the molecular events associated with the transition from ISP to DP thymocytes during the in vitro culture, RNA-seq analysis was performed on each of the four cell populations (Figure 5D). Reflecting what was observed during in vivo maturation, the in vitro maturation of WT ISP (Input_WT) to DP (cult.DP_WT) was associated with large changes in gene expression (Figure 5D; Input_WT vs cult.DP_WT). Among the 3194 genes that were differentially expressed, cell cycle related and metabolic pathway genes predominated; very few immune pathways were differentially expressed (Figure S7A and S7B). Thus, maturation of WT ISP thymocytes to the DP stage is primarily dependent on metabolic and cell cycle genes.

In the absence of BRD4, ISP cells were only able to partially progress to mature DP thymocytes, as evidenced by the lack of proliferation and low levels of CD4 expression (Figure 5A, B). About half of the genes differentially expressed between ISP and DP in the WT (1548 genes) were also differentially expressed in the BRD4-deficient cells following the in vitro culture. These common genes are mainly cell cycle related genes (Figure S7A). The genes that were differentially expressed in the transition from ISP to DP in the WT cells, but not in the BRD4-deficient cells, were mainly in metabolic pathways (Figure S7B). These results suggest that the primary defect in the maturation of ISP to DP in the absence of BRD4 may be metabolic, leading to a failure of cell cycle (Figure S7C).

The nature of the defect in maturation in the absence of BRD4 is best visualized in a principal component analysis (PCA) of the RNA-seq data (Figure 5E). As shown above (Figure 3E), the expression profiles of the WT and BRD4_cKO ISP differed significantly. During in vitro culture, the maturation of WT ISP (Input_WT) to DP thymocytes (cult.DP_WT) was accompanied by large changes in the expression profile. The BRD4_cKO ISP (Input_cKO) also underwent changes in expression profile, but were unable to fully differentiate to the DP stage (cult.DP_cKO). The CD4+ cells (cult.DP_cKO) that emerged from the overnight culture of BRD4_cKO ISP (Input_cKO) were markedly different from WT DP (cult.DP_WT) cells in their gene expression profile. Interestingly, the gene expression profile of the BRD4_cKO ISP (cult.ISP_cKO) remaining after overnight culture was more closely related to the cells that became CD4+ (cult.DP_cKO) than to the original, input ISP (Input-cKO). Thus, although the BRD4_cKO ISP can undergo some reprogramming in the DP-developmental pathway, they are unable to fully reprogram their expression to become DP thymocytes, likely due to a failure in metabolic pathways.

Taken together, our results indicate that BRD4 selectively regulates the transition from ISP to DP through its targeting of genes in metabolic pathways, as well as of myc and its targets.

### BRD4 is required for the generation of Tregs and iNKT cells, but not conventional CD4+ or CD8+ cells

Although BRD4 is required for the efficient progression of ISP to the DP stage, the DPs that develop in the absence of BRD4 further differentiate to conventional CD4+ and CD8+ single positive thymocytes (Figure 1C). This leads to the question of the effect of BRD4 loss on their gene expression profiles. Surprisingly, only a relatively small number of genes was differentially expressed in the absence of BRD4 in either CD4+ or CD8+ cells (Figure 6A, Table 1). Of those genes that were differentially expressed, the majority were positively regulated by BRD4 in both CD4+ (63%) and CD8+ (79%). This is in contrast to either the ISP or DP thymocytes where half of the genes were negatively regulated by BRD4 (Figure 3E, Table 1). Among the major pathways activated by BRD4 in both CD4+ and CD8+ are immune response pathways (Figure S8D). Although TCRβ-high single positive thymocytes can escape the BRD4 restriction and emigrate to the periphery, they are not functional and do not efficiently proliferate in response to stimulation. In addition, BRD4-deficient CD4+ and CD8+ thymocytes and splenic T cells display impaired glucose uptake, although the major metabolic pathways are not affected (Figures S8A, S8B); mitochondrial mass is reduced in the mature CD8+ cells (Figure S8C).

**Figure 6:**
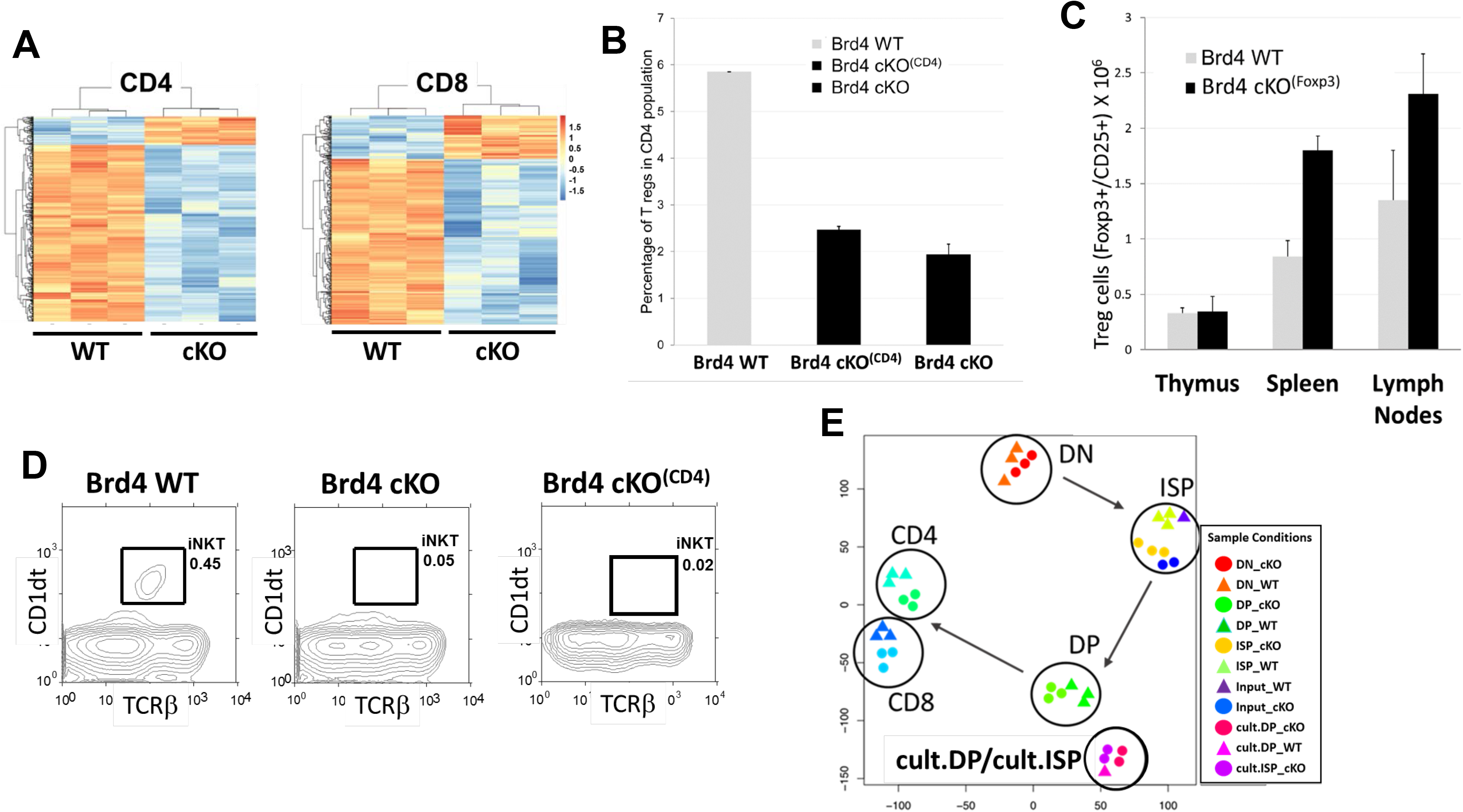
Conventional CD4+ and CD8+ single positive thymocytes develop in the absence of BRD4 but the generation of Treg and iNKT cells is affected. (A) Heat maps comparing gene expression levels in BRD4 cKO and WT CD4 and CD8 single positive thymocytes with a signal intensity fold change >2. (B) The presence of CD4+ Treg cells in BRD4-deleted and WT thymi was assessed by FACS based on internal Foxp3 expression. BRD4 deletion was mediated by either the LCK-Cre transgene (Brd4 cKO) or the CD4-Cre transgene (Brd4 cKO^(CD4)^). The graph is the summary of 3 Brd4 WT, 4 Brd4 cKO^(CD4)^ and 2 Brd4 cKO mice. (C) Quantification of CD4+ Treg cells in thymi, spleen and lymph nodes from Brd4 WT and Brd4 F/- Foxp3-Cre (cKO^(Foxp3)^) mice. (D) The presence of iNKT cells in Brd4 wild type (WT), Brd4 cKO and BRD4 cKO^(CD4)^ thymi was assessed by FACS based on CD1dt and TCRβ surface expression. The results are representative of two independent experiments. (E) T-sne representation summarizing the relationships of both in situ and in vitro thymocyte populations based on their gene expression profiles

Although conventional CD4+ and CD8+ single positive thymocytes develop in the absence of BRD4, we asked whether other thymocyte subpopulations were affected. Among the genes that were down-regulated in CD4+ thymocytes in the absence of BRD4 was Foxp3 which is required for the differentiation of Tregs (Hori et al., 2003). This led us to ask whether the development of Tregs depended on BRD4 expression. Remarkably, deletion of BRD4 by either LCK-Cre or CD4-Cre led to a significant reduction in both percentage and number of Tregs (Figure 6B) suggesting that there is a distinct requirement for BRD4 expression after the development of CD4+ and CD8+ single positive thymocytes. While BRD4 is necessary for the generation of Tregs, it is not required for maintenance of Tregs: deletion of BRD4 by Foxp3-Cre did not affect Treg development or the number of Tregs in thymus or peripheral tissues (Figure 6C). However, the ability of BRD4-deficient Tregs to proliferate in response to stimulation was markedly impaired (Figure S8E).

iNKT cell development is dependent on CD1d selection on DP thymocytes (Bendelac 1995, Zajonc 2009). BRD4 deletion did not affect CD1d1 levels in DP thymocytes (Table 2). Thus, it was surprising to find that deletion of BRD4 by either LCK-Cre or CD4-Cre resulted in a loss of iNKT cells (Figure 6D). Interestingly, BRD4 deletion by LCK-Cre resulted in a decrease in CD1d1 expression in CD4+ single positive thymocytes. This suggests that development of both iNKT and Treg subpopulations requires BRD4 expression beyond the DP stage.

Given the profound effect that BRD4 has on the differentiation of these thymocyte subpopulations, it is remarkable that its deletion targets relatively few genes. This is best visualized in a principal components analysis of all of the thymocyte subpopulations (Figure 6E) which reveals that BRD4-deleted cells still cluster closely with their WT counterpart. Notably, both the BRD4-deleted and WT ISP subpopulations are more related to one another than to either their DN precursors or DP successors, further documenting the ISP as a distinct subpopulation.

Taken together, our results demonstrate that BRD4 selectively regulates the transition from ISP to DP: it is not required either for DN differentiation or the further differentiation of conventional CD4+ and CD8+ SP thymocytes. However, it is required for the generation of Tregs and iNKT cells.

## DISCUSSION

BRD4 is being widely investigated as a therapeutic target in inflammatory diseases, as well as a variety of solid and hematological cancers (for review: (Andrieu et al., 2016; Devaiah et al., 2016b)). It is a chromatin-binding protein with both kinase and acetyl transferase activities that plays an active role in regulating transcription by linking chromatin structure and transcription (Devaiah et al., 2016a; Devaiah et al., 2012; Dey et al., 2009). In the present study, we have examined the in vivo requirement for BRD4 during thymocyte development. The stages of thymocyte differentiation are characterized by their cell surface expression of the markers CD4 and CD8: DN, ISP, DP and SP. Among these, the ISP subpopulation is the least well characterized and has been considered a transitional cell between the DN and DP stages of differentiation (Mingueneau et al., 2013). Our findings now identify ISP thymocytes as a discrete subpopulation with a gene expression profile distinct from either the DN precursors or the DP successors. Surprisingly, although BRD4 is expressed throughout all stages of thymocyte differentiation, the requirement for BRD4 is restricted to limited stages of the thymic differentiation program. Among pre-selection thymocytes, only the transition from ISP to DP is BRD4 dependent; among post-selection thymocytes, the generation of both Tregs and iNKT is abrogated in the absence of BRD4 expression. In ISP cells, BRD4 primarily regulates genes in metabolic pathways; deletion of BRD4 results in a failure to support the single round of cell cycle associated with differentiation to the DP stage. It is not required for the proliferation associated with DN stage amplification or the maturation from DP to conventional single positive thymocytes where no proliferation is involved. We propose a model in which the transition from the highly proliferative DN stage to the quiescent DP stage requires a reprogramming through the ISP stage that is regulated by BRD4.

BRD4 is expressed throughout thymocyte differentiation and at equivalent levels among the subpopulations. It is thus surprising that, among the pre-selection thymocytes, BRD4 depletion selectively affects the ISP subpopulation. The expression of over 1100 genes – most in immune, cell cycle and metabolic pathways – is affected. The loss of BRD4 results in increases in expression of about half of the genes and decreases in the other half, indicating that BRD4 acts as both a transcriptional activator and silencer. The cell cycle and metabolic pathways that are most affected by BRD4 deletion are largely down-regulated by BRD4 loss. This results in BRD4-deleted ISP that are deficient in metabolic activity.

The maturation of wild type thymocytes is accompanied by large changes in gene expression profiles. The largest changes occur in the transition from ISP to DP. Of particular note, the gene expression profile of the ISP subpopulation is distinct from both the DN and the DP; nearly 10% of its genes are uniquely differentially expressed. Thus, the ISP is not a transitional population, midway between the DN and DP. Rather, the ISPs are a molecularly and functionally distinct thymocyte subpopulation. Although the maturation from ISP to DP is accomplished during a single round of cell cycle, it is accompanied by changes in expression of approximately 40% of the genes. These differentially expressed genes map to immune, cell cycle and metabolic pathways. Among those genes are the transcription factors, TCF-1, LEF-1 and the orphan nuclear receptor, RORγt, which are necessary for the transition from ISP to DP. Significantly, their expression is not affected by BRD4 deletion and does not account for the block in differentiation of BRD4-deleted ISP. Rather, depletion of BRD4 results in the down-regulation of metabolism pathways necessary for the transition from ISP to quiescent DP cells.

Among the genes that are uniquely differentially expressed in the ISP and regulated by BRD4 is Myc. Although Myc is expressed in all thymocyte subpopulations (with the exception of DP), BRD4 only regulates Myc expression and that of its downstream targets in ISP. Surprisingly, while DN and ISP thymocytes express Myc at approximately equal levels, DN thymocytes are highly proliferative whereas ISPs undergo only a single round of proliferation. This leads to the surprising and unexpected conclusion that proliferation of ISP thymocytes is BRD4-dependent but proliferation of DN thymocytes is not. While Myc is known to be regulated by BRD4 in many, but not all, cancer cell lines, the mechanisms underlying this selective regulation of Myc by BRD4 in ISP thymocytes and other cell types remain to be determined.

In sharp contrast to the dramatic effects of BRD4 deletion on ISP thymocytes, the DN, DP, and single positive thymocyte subpopulations are only modestly affected. In the absence of BRD4, DN thymocytes differentiate normally, as evidenced by their ability to proliferate and undergo TCRβ rearrangement, consistent with the limited effect on gene expression. Similarly, BRD4 plays a relatively small role in DP or single positive thymocytes, which are phenotypically normal in the absence of BRD4. Among the genes that are down-regulated in BRD4-deficient CD4+ thymocytes is Foxp3, accounting for the failure of Tregs to develop in the absence of BRD4. Development of iNKT cells has been shown previously to be impaired by CD4-cre mediated deletion of c-myc (Dose et al., 2009). The absence of iNKT cells in thymocytes deleted of BRD4 by CD4-cre would be consistent with its regulation of Myc, except for the fact that BRD4 does not regulate cmyc in either DP or CD4 thymocytes. Equally surprising is the finding that BRD4 depletion by LCK-cre affects CD1d1 expression in CD4+ single positive thymocytes, but not DP thymocytes where it is known to be required for iNKT development. These observations suggest that iNKT development may either also require CD1d1 expression in CD4+ thymocytes or be directly regulated by BRD4, or both. Taken together, these findings lead to the unexpected conclusion that BRD4 is a major determining factor in only a subset of thymocytes, namely ISP, Treg and iNKT cells.

In conclusion, the present studies have identified the ISP as a distinct cell type that is selectively regulated by BRD4. Further maturation to the DP thymocyte stage requires a reprogramming of gene expression that depends on the BRD4-regulated pathways of Myc, cell cycle and metabolism. BRD4-independent TCR signaling and DP transcription factors are necessary, but not sufficient.

## MATERIALS AND METHODS

### Generation of the BRD4 KO mice

The constructs were described previously (Devaiah et al, 2016) The Brd4 knockout allele (brd4-) and the Brd4 floxed allele (Brd4f) were both designed to remove the third exon (carrying the ATG) of Brd4. The Brd4-allele was generated by replacing a 2kb genomic region containing exon 3 with an exogenous 6.6Kb fragment carrying the β-geo (Bgal) gene. In the Brd4f allele, exon 3 is surrounded by 2 LoxP sites. Brd4+/- mice were bred with mice expressing the Cre recombinase under the control of one of the following promoters: LCK proximal (Wilson, Taconic), CD4 (Lee et al, 2001), the Foxp3 (Chatila, 2007). Brd4f/f mice were then bred with one of the following lines, Brd4 +/- LCK-Cre**+/-**, Brd4 +/- CD4-Cre+/-, or Brd4+/- Foxp3-Cre+/- to produce Brd4F/^-^ LCK-Cre+/- or Brd4F/^-^ CD4-Cre+/- or Brd4F/^-^ Foxp3-Cre+/- mice, respectively. The Brd4 f/f vav-cre mice were generated by crossing the Brd4 f/f mice with mice expressing the cre recombinase under the vav promoter (Georgiades et al 2002). Cells were prepared from thymus, counted and assessed for CD4, CD8 and TCRβ surface protein expression by flow cytometry using either FACS Aria or Fortessa.

### Purification of DN thymocytes and lymph node CD4 and CD8 T cells

DP thymocytes were purified by flow cytometry based on the expression of both CD4 and CD8 surface markers and the lack of CD69 at the surface. CD4 thymocytes were pre-purified using the Life Technology Dynabeads Untouched Mouse CD4 Cells Kit and sorted by flow cytometry based on the expression of CD4 and TCRβ at the surface. DN, ISP and CD8 thymocytes were pre-purified using the Life Technology Dynabeads Untouched Mouse CD8 Cells Kit and then sorted by flow cytometry respectively on their lack of CD4, CD8, TCRβ surface expression for DN, the expression of CD8 and the lack of TCRβ expression for the ISP and finally the expression of CD8 and TCRβ at the surface for CD8 Single Positive cells. Lymph node CD4 and CD8 T cells were purified using respectively the EasySep Mouse CD4+ or CD8+ T cell Isolation kit (StemCell Technologies) followed by FACS based on CD4, CD8 and TCRβ expression.

### Antibodies

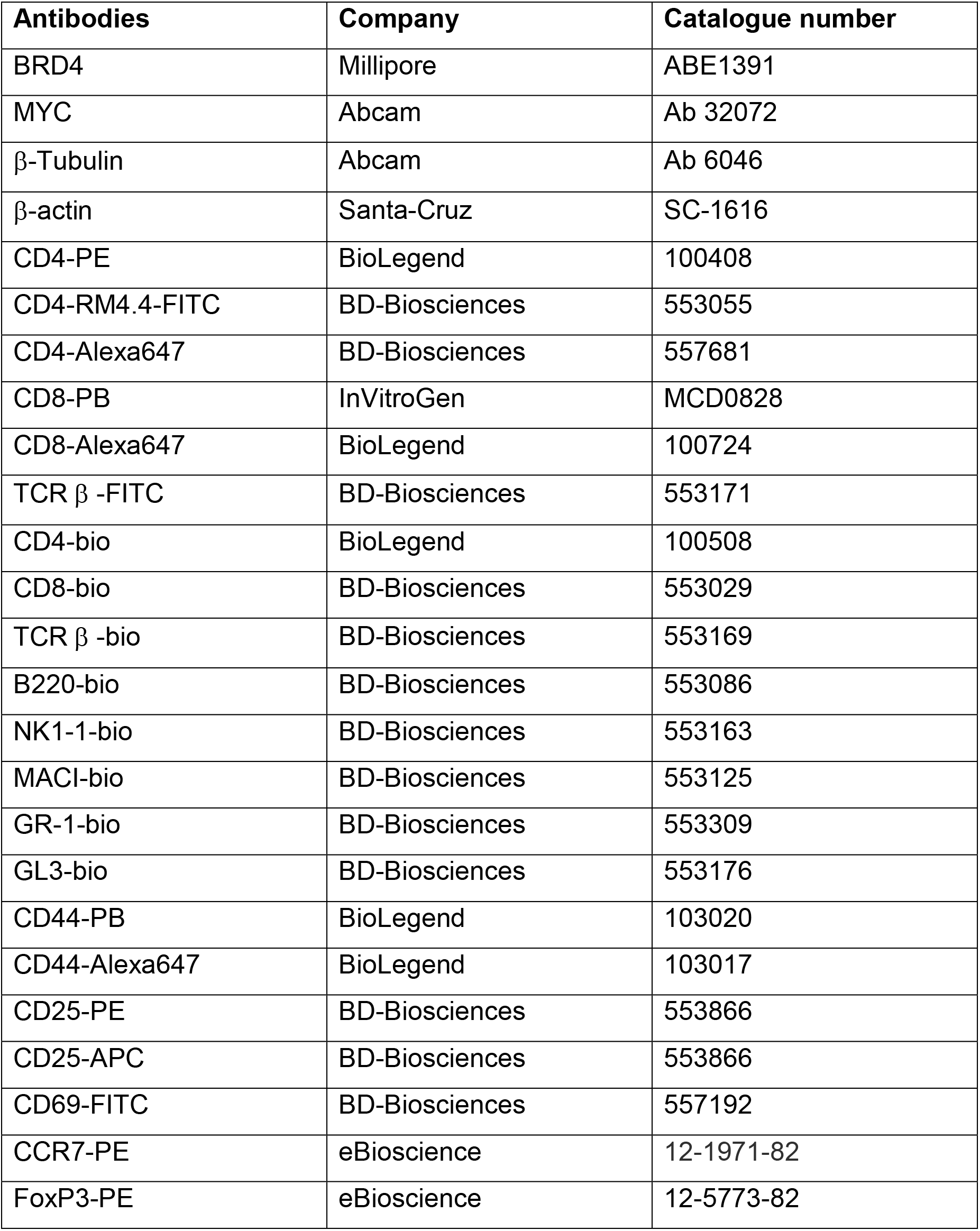

**Antibodies.**
| **Antibodies** | **Company** | **Catalogue number** |
| --- | --- | --- |
| BRD4 | Millipore | ABE1391 |
| MYC | Abcam | Ab 32072 |
| β-Tubulin | Abcam | Ab 6046 |
| β-actin | Santa-Cruz | SC-1616 |
| CD4-PE | BioLegend | 100408 |
| CD4-RM4.4-FITC | BD-Biosciences | 553055 |
| CD4-Alexa647 | BD-Biosciences | 557681 |
| CD8-PB | InVitroGen | MCD0828 |
| CD8-Alexa647 | BioLegend | 100724 |
| TCR β -FITC | BD-Biosciences | 553171 |
| CD4-bio | BioLegend | 100508 |
| CD8-bio | BD-Biosciences | 553029 |
| TCR β -bio | BD-Biosciences | 553169 |
| B220-bio | BD-Biosciences | 553086 |
| NK1-1-bio | BD-Biosciences | 553163 |
| MAC1-bio | BD-Biosciences | 553125 |
| GR-1-bio | BD-Biosciences | 553309 |
| GL3-bio | BD-Biosciences | 553176 |
| CD44-PB | BioLegend | 103020 |
| CD44-Alexa647 | BioLegend | 103017 |
| CD25-PE | BD-Biosciences | 553866 |
| CD25-APC | BD-Biosciences | 553866 |
| CD69-FITC | BD-Biosciences | 557192 |
| CCR7-PE | eBioscience | 12-1971-82 |
| FoxP3-PE | eBioscience | 12-5773-82 |

**Western blotting**. 5 X 10^5^ either thymocytes, or lymph node lympocytes were extracted using the TER1 buffer (ThermoFisher, FNN0071) and the soluble lysate fractions analysed by western blotting using the BRD4 antibody, kindly provided by Dr Ozato. BRD4 short form was detected using Millipore BRD4 antibody (ABE1391) and Myc using the Abcam Y69 (Ab 32072). Tubulin(), and β-actin antibodies were used as loading contriol.

### RNA-seq analysis

RNA-seq analysis was done as described before^(ref)^. RNAs were isolated from WT and BRD4 deleted cells using the microRNeasy-plus kit (Qiagen). Libraries were made using the Clontech SMARTer Universal Low Input RNA Kit for Sequencing and sequenced as pair-end reads on an Hiseq platform according to established procedures. RNA-seq reads were aligned to mouse reference genome (mm10) with STAR aligner (1). Raw read counts were obtained using htseq-count (2) and normalized for further analysis using the built-in normalization algorithms of DESeq2 (3). All differential expression analysis was performed with DESeq2 and a gene was considered as differentially expressed between two groups when its fold-change > 2 and FDR adjusted P-value < 0.01. Statistical testing, principal component analysis (PCA), hierarchical clustering and t-SNE(4) were performed in R. Gene set enrichment analysis (GSEA) for the differentially expressed genes was performed using Hallmark gene set collection from MsigDB (http://www.broad.mit.edu/gsea/).

Read visualization was assessed using the IGV platform. GO and Pathway analysis were also addressed using Metascape.org.

#### BrdU incorporation

Mice were injected in the peritoneal cavity with 1mg of BrdU twice, two and four hours before harvesting the thymi. The DN thymocytes were stained using a cocktail of antibodies recognizing surface proteins expressed on lineage committed cells (TCRβ, CD4, CD8, B220, NK1-1, MACI, GR-1, GL3) and CD44 and CD25 antibodies; ISP thymocytes were stained with the samecocktail of antibodies minus CD8. and then fixed and treated according to the procedure described in the BrdU flow kit (BD Pharmingen, 559619). DN thymocytes were identified by the absence of staining with a cocktail of antibodies recognizing surface proteins expressed on lineage committed cells (TCRβ, CD4, CD8, B220, NK1-1, MACI, GR-1), the subpopulations of DN and ISP were defined based on the expression of CD44 and CD25 or CD8, respectively, and their ability to proliferate assessed by flow cytometry based on BrdU incorporation.

#### TCRβ intracellular staining

DN thymocytes were identified as above, fixed using the cytofix/cytoperm solution and the procedure recommended by BD Pharmingen, internally stained with an anti TCRβ antibody and analysed by flow cytometry.

### Glucose uptake

2 10^6^ cells were washed once with serum free RPMI then resuspended in 1ml of warmed serum free RPMI. Mitotracker Green (ThermoFisher M7514) was diluted to a 1uM working solution, then added to the cells at the final concentration of 100nM. Cells were incubated for 20 mn at 37oC, then washed once prior to staining with surface markers.

### Mitochondrial mass evaluation

2 10^6^ cells were washed once with serum free/ glucose free RPMI then resuspended in 1ml of warmed serum free/ glucose free RPMI. 2-NBDG (ThermoFisher, N 13195) was diluted to a 1mM working solution in DMSO, then added to the cells at the final concentration of 100uM. Cells were incubated for 20 mn at 37oC, then washed once prior to staining with surface markers.

### ISP culture

Either WT or Brd4 deleted ISP Thymocytes were purified as described above using the Life Technology Dynabeads Untouched Mouse CD8 Cells Kit and then sorted by FACS based on their CD8 surface expression and the lack of TCRβ surface expression. Equal amount of WT and Brd4 deleted cells were grown for 16h in RPMI medium supplemented with 10% fetal calf serum, Na Pyruvate 1mM, MEM non essential Amino Acids 1X and 2-mercaptoethanol and their ability to proliferate assayed by counting the cells after culture. The cells were also analysed by Flow cytometry for their ability to express CD4 at their surface.

### Tregs analysis

Treg thymocytes were fixed using the Foxp3/TranscriptionFactor Staining buffer kit from eBioscience (00–5523–00), internally stained with an anti Foxp3 antibody and analysed by flow cytometry. For anti-TCR–induced T-cell proliferation, Treg cells (3 to 5 × 10^4^/well) were placed in 96-well round-bottom plates (0.2 mL) together with irradiated T cell–depleted B6 spleen cells (2000R) as accessory cells (APCs) and stimulated with anti-CD3 mAb (1 μg/mL) with or without rIL-2 (200 U/mL) for 72 hours. Cultures were pulsed with [^3^H]-thymidine 8 hours before harvest.

## ONLINE SUPPLEMENTAL MATERIAL

Figure S1 shows the relative abundance of the BRD4 proteins in each thymocyte population and splenic lymphocytes. Figure S2A represents the schema of the Floxed allele; Figure S2B, the T cell development steps and Figure S2C, extent of Brd4 deletion as assessed by RNAseq analysis. Figure S3 shows the effect of BRD4 deletion on expression of different genes that characterize immature ISP. Figure S4 shows the decrease in the size of DN4 and ISP thymocytes that accompanies the loss of BRD4. Figure S5 summarizes the pathways enriched in WT ISP relative to DN or/and DP and Figure S6, the pathways affected by BRD4 deletion in ISP and DP thymocytes. Figure S7 shows the pathways that are affected as WT and cKO ISP mature in cutlure. Figures S8A, S8B, S8C, S8D summarize the effect of BRD4 deletion on CD4 and CD8 thymocytes, Figure S8E compares the ability of BRD4 deleted and WT Tregs to proliferate.

## AUTHOR CONTRIBUTIONS

A.G. conceived, designed and executed the experiments and wrote the manuscript; Q-R.C. and D.M. performed the analysis of the RNA-seq data; A.D, and K.O. provided the BRD4fl/fl and BRD4+/- mice; R.E. and X.T. helped with the in vitro cultures of ISP and Tregs, respectively; A.S. and D.S. conceptualized and interpreted the data and wrote the manuscript

## ACKNOWLEDGEMENTS

The authors gratefully acknowledge Drs. Remy Bosselut, Richard Hodes, Hyun Park, and Ranjan Sen for their critical reading of the manuscript and helpful discussions. Dr. Jie Mu and Dr. Thomas Ciucci for their technical assistance and data analysis support, respectively. We also thank members of the D. Singer lab for discussions. This research was supported by the Intramural Research Program of the NIH, National Cancer Institute Center for Cancer Research.

## REFERENCES

Alsarraj, J., Faraji, F., Geiger, T.R., Mattaini, K.R., Williams, M., Wu, J., Ha, N.H., Merlino, T., Walker, R.C., Bosley, A.D., et al. (2013). BRD4 short isoform interacts with RRP1B, SIPA1 and components of the LINC complex at the inner face of the nuclear membrane. PLoS One 8, e80746.

Andrieu, G., Belkina, A.C., and Denis, G.V. (2016). Clinical trials for BET inhibitors run ahead of the science. Drug discovery today. Technologies 19, 45–50.

Barrow, J.J., Balsa, E., Verdeguer, F., Tavares, C.D., Soustek, M.S., Hollingsworth, L.R.t., Jedrychowski, M., Vogel, R., Paulo, J.A., Smeitink, J., et al. (2016). Bromodomain Inhibitors Correct Bioenergetic Deficiency Caused by Mitochondrial Disease Complex I Mutations. Mol Cell 64, 163–175.

Bolden, J.E., Tasdemir, N., Dow, L.E., van Es, J.H., Wilkinson, J.E., Zhao, Z., Clevers, H., and Lowe, S.W. (2014). Inducible in vivo silencing of Brd4 identifies potential toxicities of sustained BET protein inhibition. Cell Rep 8, 1919–1929.

Brown, J.D., Lin, C.Y., Duan, Q., Griffin, G., Federation, A.J., Paranal, R.M., Bair, S., Newton, G., Lichtman, A.H., Kung, A.L., et al. (2014). NF-kappaB directs dynamic super enhancer formation in inflammation and atherogenesis. Mol Cell 56, 219–231.

Cheung, K.L., Zhang, F., Jaganathan, A., Sharma, R., Zhang, Q., Konuma, T., Shen, T., Lee, J.Y., Ren, C., Chen, C.H., et al. (2017). Distinct Roles of Brd2 and Brd4 in Potentiating the Transcriptional Program for Th17 Cell Differentiation. Mol Cell 65, 1068–1080.e1065.

Deeney, J.T., Belkina, A.C., Shirihai, O.S., Corkey, B.E., and Denis, G.V. (2016). BET Bromodomain Proteins Brd2, Brd3 and Brd4 Selectively Regulate Metabolic Pathways in the Pancreatic beta-Cell. PLoS One 11, e0151329.

Devaiah, B.N., Case-Borden, C., Gegonne, A., Hsu, C.H., Chen, Q., Meerzaman, D., Dey, A., Ozato, K., and Singer, D.S. (2016a). BRD4 is a histone acetyltransferase that evicts nucleosomes from chromatin. Nat Struct Mol Biol.

Devaiah, B.N., Gegonne, A., and Singer, D.S. (2016b). Bromodomain 4: a cellular Swiss army knife. J Leukoc Biol 100, 679–686.

Devaiah, B.N., Lewis, B.A., Cherman, N., Hewitt, M.C., Albrecht, B.K., Robey, P.G., Ozato, K., Sims, R.J., 3rd, and Singer, D.S. (2012). BRD4 is an atypical kinase that 28 phosphorylates serine2 of the RNA polymerase II carboxy-terminal domain. Proc Natl Acad Sci U S A 109, 6927–6932.

Dey, A., Ellenberg, J., Farina, A., Coleman, A.E., Maruyama, T., Sciortino, S., Lippincott-Schwartz, J., and Ozato, K. (2000). A bromodomain protein, MCAP, associates with mitotic chromosomes and affects G(2)-to-M transition. Mol Cell Biol 20, 6537–6549.

Dey, A., Nishiyama, A., Karpova, T., McNally, J., and Ozato, K. (2009). Brd4 marks select genes on mitotic chromatin and directs postmitotic transcription. Mol Biol Cell 20, 4899–4909.

Di Micco, R., Fontanals-Cirera, B., Low, V., Ntziachristos, P., Yuen, S.K., Lovell, C.D., Dolgalev, I., Yonekubo, Y., Zhang, G., Rusinova, E., et al. (2014). Control of embryonic stem cell identity by BRD4-dependent transcriptional elongation of super-enhancer-associated pluripotency genes. Cell Rep 9, 234–247.

Dose, M., Sleckman, B.P., Han, J., Bredemeyer, A.L., Bendelac, A., and Gounari, F. (2009). Intrathymic proliferation wave essential for Valpha14+ natural killer T cell development depends on c-Myc. Proc Natl Acad Sci U S A 106, 8641–8646.

Farina, A., Hattori, M., Qin, J., Nakatani, Y., Minato, N., and Ozato, K. (2004). Bromodomain protein Brd4 binds to GTPase-activating SPA-1, modulating its activity and subcellular localization. Mol Cell Biol 24, 9059–9069.

Georgiades, P., Ogilvy, S., Duval, H., Licence, D.R., Charnock-Jones, D.S., Smith, S.K., and Print, C.G. (2002). VavCre transgenic mice: a tool for mutagenesis in hematopoietic and endothelial lineages. Genesis (New York, N.Y.: 2000) 34, 251–256.

Hori, S., Nomura, T., and Sakaguchi, S. (2003). Control of regulatory T cell development by the transcription factor Foxp3. Science (New York, N.Y.) 299, 1057–1061.

Houzelstein, D., Bullock, S.L., Lynch, D.E., Grigorieva, E.F., Wilson, V.A., and Beddington, R.S. (2002). Growth and early postimplantation defects in mice deficient for the bromodomain-containing protein Brd4. Mol Cell Biol 22, 3794–3802.

Jang, M.K., Mochizuki, K., Zhou, M., Jeong, H.S., Brady, J.N., and Ozato, K. (2005). The bromodomain protein Brd4 is a positive regulatory component of P-TEFb and stimulates RNA polymerase II-dependent transcription. Mol Cell 19, 523–534.

Lee, P.P., Fitzpatrick, D.R., Beard, C., Jessup, H.K., Lehar, S., Makar, K.W., Perez-Melgosa, M., Sweetser, M.T., Schlissel, M.S., Nguyen, S., et al. (2001). A critical role for Dnmt1 and DNA methylation in T cell development, function, and survival. Immunity 15, 763–774.

Loven, J., Hoke, H.A., Lin, C.Y., Lau, A., Orlando, D.A., Vakoc, C.R., Bradner, J.E., Lee, T.I., and Young, R.A. (2013). Selective inhibition of tumor oncogenes by disruption of super-enhancers. Cell 153, 320–334.

Maruyama, T., Farina, A., Dey, A., Cheong, J., Bermudez, V.P., Tamura, T., Sciortino, S., Shuman, J., Hurwitz, J., and Ozato, K. (2002). A Mammalian bromodomain protein, brd4, interacts with replication factor C and inhibits progression to S phase. Mol Cell Biol 22, 6509–6520.

Mele, D.A., Salmeron, A., Ghosh, S., Huang, H.R., Bryant, B.M., and Lora, J.M. (2013). BET bromodomain inhibition suppresses TH17-mediated pathology. J Exp Med 210, 2181–2190.

Mingueneau, M., Kreslavsky, T., Gray, D., Heng, T., Cruse, R., Ericson, J., Bendall, S., Spitzer, M.H., Nolan, G.P., Kobayashi, K., et al. (2013). The transcriptional landscape of alphabeta T cell differentiation. Nat Immunol 14, 619–632.

Mochizuki, K., Nishiyama, A., Jang, M.K., Dey, A., Ghosh, A., Tamura, T., Natsume, H., Yao, H., and Ozato, K. (2008). The bromodomain protein Brd4 stimulates G1 gene transcription and promotes progression to S phase. J Biol Chem 283, 9040–9048.

Roberts, T.C., Etxaniz, U., Dall’Agnese, A., Wu, S.Y., Chiang, C.M., Brennan, P.E., Wood, M.J.A., and Puri, P.L. (2017). BRD3 and BRD4 BET Bromodomain Proteins Differentially Regulate Skeletal Myogenesis. Scientific reports 7, 6153.

Rothenberg, E.V., Ungerback, J., and Champhekar, A. (2016). Forging T-Lymphocyte Identity: Intersecting Networks of Transcriptional Control. Adv Immunol 129, 109–174.

Sakamaki, J.I., Wilkinson, S., Hahn, M., Tasdemir, N., O’Prey, J., Clark, W., Hedley, A., Nixon, C., Long, J.S., New, M., et al. (2017). Bromodomain Protein BRD4 Is a Transcriptional Repressor of Autophagy and Lysosomal Function. Mol Cell 66, 517–532.e519.

Schmidt, S.F., Larsen, B.D., Loft, A., Nielsen, R., Madsen, J.G., and Mandrup, S. (2015). Acute TNF-induced repression of cell identity genes is mediated by NFkappaB-directed redistribution of cofactors from super-enhancers. Genome Res 25, 1281–1294.

Takahama, Y., Shores, E.W., and Singer, A. (1992). Negative selection of precursor thymocytes before their differentiation into CD4+CD8+ cells. Science (New York, N.Y.) 258, 653–656.

Tasdemir, N., Banito, A., Roe, J.S., Alonso-Curbelo, D., Camiolo, M., Tschaharganeh, D.F., Huang, C.H., Aksoy, O., Bolden, J.E., Chen, C.C., et al. (2016). BRD4 Connects Enhancer Remodeling to Senescence Immune Surveillance. Cancer Discov.

Vacchio, M.S., Ciucci, T., and Bosselut, R. (2016). 200 Million Thymocytes and I: A Beginner’s Survival Guide to T Cell Development. Methods in molecular biology (Clifton, N.J.) 1323, 3–21.

Whyte, W.A., Orlando, D.A., Hnisz, D., Abraham, B.J., Lin, C.Y., Kagey, M.H., Rahl, P.B., Lee, T.I., and Young, R.A. (2013). Master transcription factors and mediator establish super-enhancers at key cell identity genes. Cell 153, 307–319.

Wu, T., Pinto, H.B., Kamikawa, Y.F., and Donohoe, M.E. (2015). The BET family member BRD4 interacts with OCT4 and regulates pluripotency gene expression. Stem Cell Reports 4, 390–403.

Yang, Z., Yik, J.H., Chen, R., He, N., Jang, M.K., Ozato, K., and Zhou, Q. (2005). Recruitment of P-TEFb for stimulation of transcriptional elongation by the bromodomain protein Brd4. Mol Cell 19, 535–545.

You, J., Croyle, J.L., Nishimura, A., Ozato, K., and Howley, P.M. (2004). Interaction of the bovine papillomavirus E2 protein with Brd4 tethers the viral DNA to host mitotic chromosomes. Cell 117, 349–360.

You, J., Li, Q., Wu, C., Kim, J., Ottinger, M., and Howley, P.M. (2009). Regulation of aurora B expression by the bromodomain protein Brd4. Mol Cell Biol 29, 5094–5103.

You, J., Schweiger, M.R., and Howley, P.M. (2005). Inhibition of E2 binding to Brd4 enhances viral genome loss and phenotypic reversion of bovine papillomavirus-transformed cells. J Virol 79, 14956–14961.

Yu, Q., Erman, B., Park, J.H., Feigenbaum, L., and Singer, A. (2004). IL-7 receptor signals inhibit expression of transcription factors TCF-1, LEF-1, and RORgammat: impact on thymocyte development. J Exp Med 200, 797–803.

Zuber, J., Shi, J., Wang, E., Rappaport, A.R., Herrmann, H., Sison, E.A., Magoon, D., Qi, J., Blatt, K., Wunderlich, M., et al. (2011). RNAi screen identifies Brd4 as a therapeutic target in acute myeloid leukaemia. Nature 478, 524–528.

